# Chaperoning through time - defining DNAJA2 co-chaperone client selectivity via pulse-SILAC and BioID

**DOI:** 10.64898/2026.07.16.739051

**Authors:** Shriya Kamat, Gaetano Calabrese, Vanessa Emnacen-Pankhurst, Chloe Shacham Dupont, Elizabeth Hui, Hailey Eng, Sahil Chandhok, Thibault Mayor

## Abstract

Molecular chaperones are a major driving force ensuring efficient protein folding and regulating proteostasis in the cell. However, it remains unclear how their clients are selected in most cases, especially after the release of nascent protein chains from ribosomes. Here, we present a novel approach that combines pulse metabolic labelling with SILAC and BioID mass spectrometry to temporally resolve protein-protein interactions of a co-chaperone. Using the Hsp70 co-chaperone DNAJA2 as a benchmark, we reveal that two distinct pools of proteins are enriched. In particular, DNAJA2 displays a preferential association with highly structured recently synthesized proteins enriched in β-strands. In contrast, pre-existing proteins captured in our assay exhibit higher intrinsic disorder. In both cases, these proteins tend to be longer and contain a lower net charge compared to the proteome. Notably, these preferential associations are retained upon heat-shock, while interactions with the “older” pool of proteins become more prevalent under these conditions. Through this methodology, we gain novel insights on how co-chaperone-client interactions may occur over the lifespan of a protein to preserve proteostasis.

## INTRODUCTION

Protein functionality is typically dependent on maintaining a native state by folding into a defined three-dimensional structure, a process often challenged by molecular crowding and a relatively slow folding rate in cells^1–4^. Pertubations to protein homeostasis (proteostasis), arising from factors such as genetic mutations, oxidative stress, or aging, can further impair this process and lead to protein misfolding and aggregation, a hallmark of numerous proteinopathies including neurodenerative diseases such as Alzheimer’s and Huntington’s diseases ^3–12^. These challenges underscore the need for a coordinated proteostasis network to ensure efficient protein folding and prevent aggregation. Notably, molecular chaperones are key players evolved to ensure protein folding occurs efficiently in a biologically-relevant time scale^9,13^.

Protein folding begins during translation^14^ and the integration of ribosome profiling with chaperone enrichment strategies has provided detailed insight onto where molecular chaperones engage nascent polypeptide chains cotranslationally^15,16^. For instance, the ribosome-associated chaperones such as the nascent-polypeptide associated complex (NAC) engage nascent chains at the ribosomal exit tunnel and prevent misfolding of emerging polypeptides^14,17–19^. The 70kDa Heat shock protein (Hsp70) chaperones that recognize and bind hydrophobic stretches of amino acids are further recruited, to ensure off-pathway aggregation is not pursued^20–22^. As elongation proceeds, recruitment of downstream chaperone systems, notably the TRiC/CCT chaperonin, also occurs to ensure proteins attain their native states^23,24^. Nonetheless, our recent pulse-SILAC study revealed that a subset of proteins remains thermodynamically less stable immediately following synthesis, indicating that the acquisition of native-state stability is not always completed cotranslationally^25^. These proteins are generally abundant and structurally complex, highlighting an increased dependence on post-translational protein homeostasis mechanisms for their maturation.

Molecular chaperones continue to govern the polypeptide’s fate beyond *de novo* folding^9,26^. For instance, the Hsp70/40 system plays central roles in post-translational folding, trafficking, complex assembly, disagreggation and protein quality control. Through ATP-dependent cycles of client binding and release, Hsp70s maintain unfolded or partially folded proteins in a soluble, folding-competent state, and promote refolding^26^. This stabilization also facilitates interactions with targeting machineries that direct folding intermediates to specific cellular compartments or complex formation^27^. They also promote disagregation^28–32^ or can mediate the triage towards ubiqutin-dependent proteasome or lysosomal degradation^33,34^.

The versatality of the Hsp70s is largely enabled by their co-chaperones, Hsp40/DnaJ proteins^35,36^. DnaJs stimulate the Hsp70’s chaperone activity through the conserved J-domain inducing a conformational change to promote client capture^37^. In humans, there are 49 DnaJ proteins divided in three A, B and C classes^13,35,36^ and some of these DnaJs appear to possess functions extending beyond their canonical roles as Hsp70 co-chaperones, further highlighting the complexity^38,39^. Several proteomic studies have began to map the interactome of eukaryotic DnaJ chaperones and their cognate Hsp70s to gain insight into their specific function and client repertoires^40–42^; Particularly, approaches involving proximity-labelling (BioID-MS) have proven particularly useful for capturing potential transient and low-affinity co-chaperone-client interactions *in vivo*^41,43,44^. Despite these efforts, a larger gap remains in understanding how DnaJs select and engage proteins across different stages of their lifecycle, from nascent chains to post-translational maintanance and eventual degradation.

In our study, we combined pulse-SILAC with BioID-MS to better define the temporal relation of interacting proteins by leveraging metabolic labelling to discriminate recently synthesized from pre-existing protein populations. Notably, we show that DNAJA2 interactions are preferentially enriched for highly structured recently synthesized proteins and disordered, pre-existing proteins in both stressed and unstressed conditions, providing insight into its role in maintaining proteostasis.

## RESULTS

### Establishing a pulse-SILAC with BioID-MS methodology in mammalian HEK293 cells

We sought to establish a method to temporally resolve co-chaperone-client interactions by combining pulse-SILAC with BioID mass spectrometry. Briefly, HEK293 cells expressing tetracycline-inducible DNAJA1 and DNAJA2 C-terminally tagged with FLAG-BioID2 were pulse labelled with heavy SILAC media (^13^C/^15^N-containing lysine and arginine residues) together with biotin for two hours prior to cell lysis (Fig. 1A, 1B). We verified that the two co-chaperone baits and the control (Renilla luciferase) localized predominantly in the cytosol and that biotinylation signals were above background in these conditions (Fig. S1A, S1B). Cell lysates were collected, and a portion of the total cell lysate (TCL) was directly analyzed by mass spectrometry. The remainder of the samples was subjected to the BioID pulldown workflow using streptavidin affinity purification to isolate biotinylated proteins for mass spectrometry analysis.

**Figure 1.**
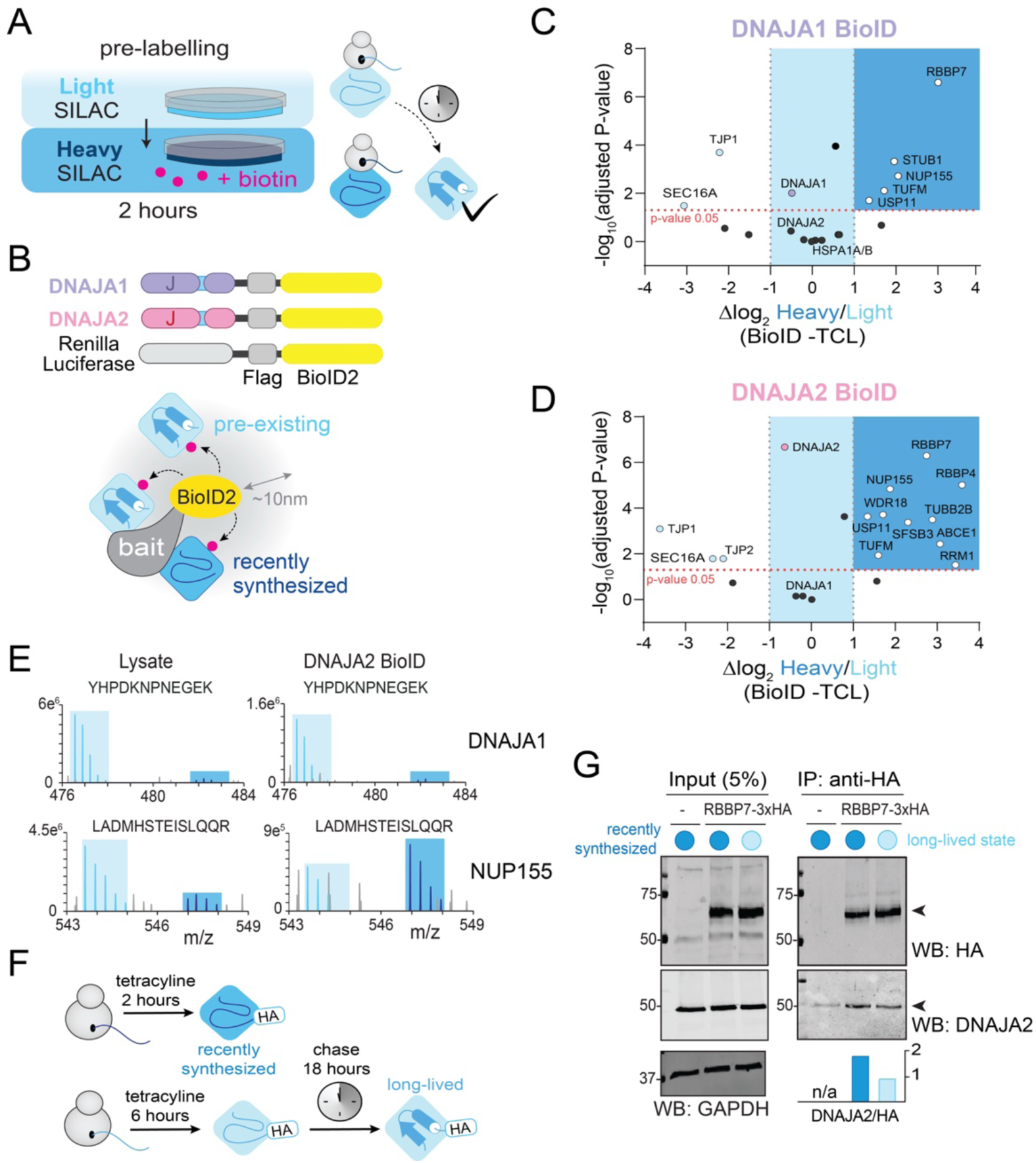
Pulse-SILAC combined with bioID in mammalian tissue culture cells. **A and B.** Schematic representation of a single pulse-SILAC experiment (A) combined with BioID labelling (B). Cells are pulsed with heavy SILAC medium for two hours to distinguish recently synthesized proteins (dark blue) from pre-existing proteins that have had more time to mature (light blue). Because biotin is added only during the pulse-SILAC labelling period, recently synthesised that are biotinylated due to their proximity from the bait are expected to exhibit a strong heavy SILAC incorporation, whereas pre-existing proteins should show little or no heavy labelling. **C and D.** Volcano plots showing the log_2_ H/L ratios of high confidence interactors (BFDR<0.05) of DNAJA1 (C) and DNAJA2 (D) quantified in the BioID samples and normalized to SILAC ratios in the TCL samples. Statistical significance (p-adj <0.05) calculated using an unpaired t-test adjusted using a Holm-Šídák based correction. **E.** Representative MS1 spectra of the indicated NUP155 and DNAJA1 peptides from the BioID and the TCL fractions of the DNAJA2 experiment. **F.** Schematic of the two conditions used for the reverse co-IP experiment to enrich either recently synthesized RBB7-3xHA (dark blue) or long-lived species (light blue). **G.** Western blot analysis of reverse co-IP using anti-HA magnetic beads to pull down RBBP7-3xHA that was expressed for 2 hours or for 6 hours followed by an 18-hour chase. HEK293T T-REx Flp-In cells were used as negative controls. Quantification of the DNAJA2 co-precipitation signal was performed by subtracting the background signal from the WT control and normalizing to HA signal.

In this pilot experiment 2544 and 2577 proteins were quantified in the TCL fractions of the DNAJA1 and DNAJA2 samples, respectively (Table S1). Across all samples, ∼17% of the protein pool was heavy SILAC-labelled, based on the median SILAC ratios of each sample, indicating that a sizeable portion of the analyzed proteome consisted of recently synthesized proteins. We next identified high-confidence interactors for each bait by first summing the SILAC intensity signals of each potential prey, and then compared their heavy-to-light (H/L) SILAC ratios relative to those observed in the corresponding TCL samples. Proteins exhibiting significantly higher H/L SILAC ratios (log_2_ of fold change (FC) > 1) were classified as “recently-synthesized” candidate-clients, as these likely represent proteins engaging with the co-chaperone co-translationally or post-translationally during the two-hour heavy SILAC labelling period.

Out of the 42 candidate interactors identified for DNAJA1 (BFDR <0.05), 19 were also quantified in the TCL fraction, including six recently-synthesized candidate clients (Fig. 1C). For comparison, the DNAJA1 bait itself displayed no shift in SILAC labelling, similar to its interacting proteins HSPA1A and DNAJA2. In these cases, the chaperone and co-chaperone are presumably mature proteins that associate with DNAJA1 in the cell. Therefore, proteins with log_2_ H/L SILAC ratios between −1 and 1 likely represent interacting proteins with no particular enrichment for recently synthesized species. These may include proteins functioning together with the co-chaperone or clients that continue to interact with the chaperone throughout their lifespan. Interestingly, proteins such as the tight junction protein TJP1 and the ER-associated protein SEC16A displayed a significant depletion of recently synthesized species, suggesting that the co-chaperone may preferentially interact with older pools of these proteins.

Similarly, 31 high-confidence candidate interactors were identified for DNAJA2, of which 19 were quantified in the TCL fraction, including 10 recently synthesized proteins (Fig. 1D). As before, the SILAC ratios of the bait and the interacting co-chaperone DNAJA1 remained unchanged, whereas the HSPA1A did not pass the SAINT statistical threshold. We were also unable to identify many high-confidence interactors when using HSPA1A as a bait under these conditions, despite strong biotinylation signal (Fig. S1, S2, Table S1). Notably, the nucleoporin NUP155 was identified as a recently synthesized candidate client for both co-chaperones. For example, the LADMHSTEISLQQR peptide from NUP155 displayed a clear shift in SILAC labelling between the TCL and streptavidin-enriched fractions following labelling with DNAJA2-BioID2 (Fig. 1E). In contrast, the YHPDKNPNEGEK peptide derived from DNAJA1 in the same experiment displayed no apparent change in SILAC distribution. These results indicate that proteins biotinylated in presence of the two co-chaperones have been synthesized at different times.

To validate this preferential interaction with recently synthesized proteins, we used an orthogonal approach based on differential expressions. We selected the core-histone binding protein RBBP7, a candidate interactor identified in both experiments that was fused to HA tag and expressed either for two hours following tetracycline induction (to mimic recently synthesized conditions) or for 6 hours followed by an 18-hour chase (to mimic pre-existing conditions) (Fig. 1F). Following chemical cross-linking to stabilize potential weak interactions, endogenous DNAJA2 was co-immunoprecipitated at higher levels with the recently synthesized RBBP7 with anti-HA beads (Fig. 1G). Approximately twofold higher levels of DNAJA2 were recovered in the recently synthesized condition relative to the long-lived condition, supporting the observations from the mass spectrometry experiments. Together, these results demonstrate that combining pulse-SILAC with BioID enables temporal resolution of co-chaperone-client interactions in mammalian tissue culture cells.

### Distinguishing DNAJA2’s temporal interactome via double-pulse SILAC

Our pilot experiment indicates that certain proteins preferably interact with the two co-chaperones up to two hours post synthesis. We wondered whether the interaction with the co-chaperone could persist beyond two hours indicative of a relatively slow protein maturation process. Therefore, we devised a double pulse-SILAC experiment (Fig. 2A). We opted to benchmark the methodology using DNAJA2, as this co-chaperone was observed to have a slightly higher number of candidate interactors in the pilot experiment. HEK293 cells expressing tetracycline-inducible DNAJA2-FLAG-BioID2 were first prelabeled with heavy SILAC, then pulsed with medium SILAC for two hours before switching for a final two-hour incubation with light SILAC containing biotin (Fig. 2A). In this particular case, we selected to swap the labelling order for additional control (i.e., the pool of recently synthesized proteins is light SILAC).

**Figure 2.**
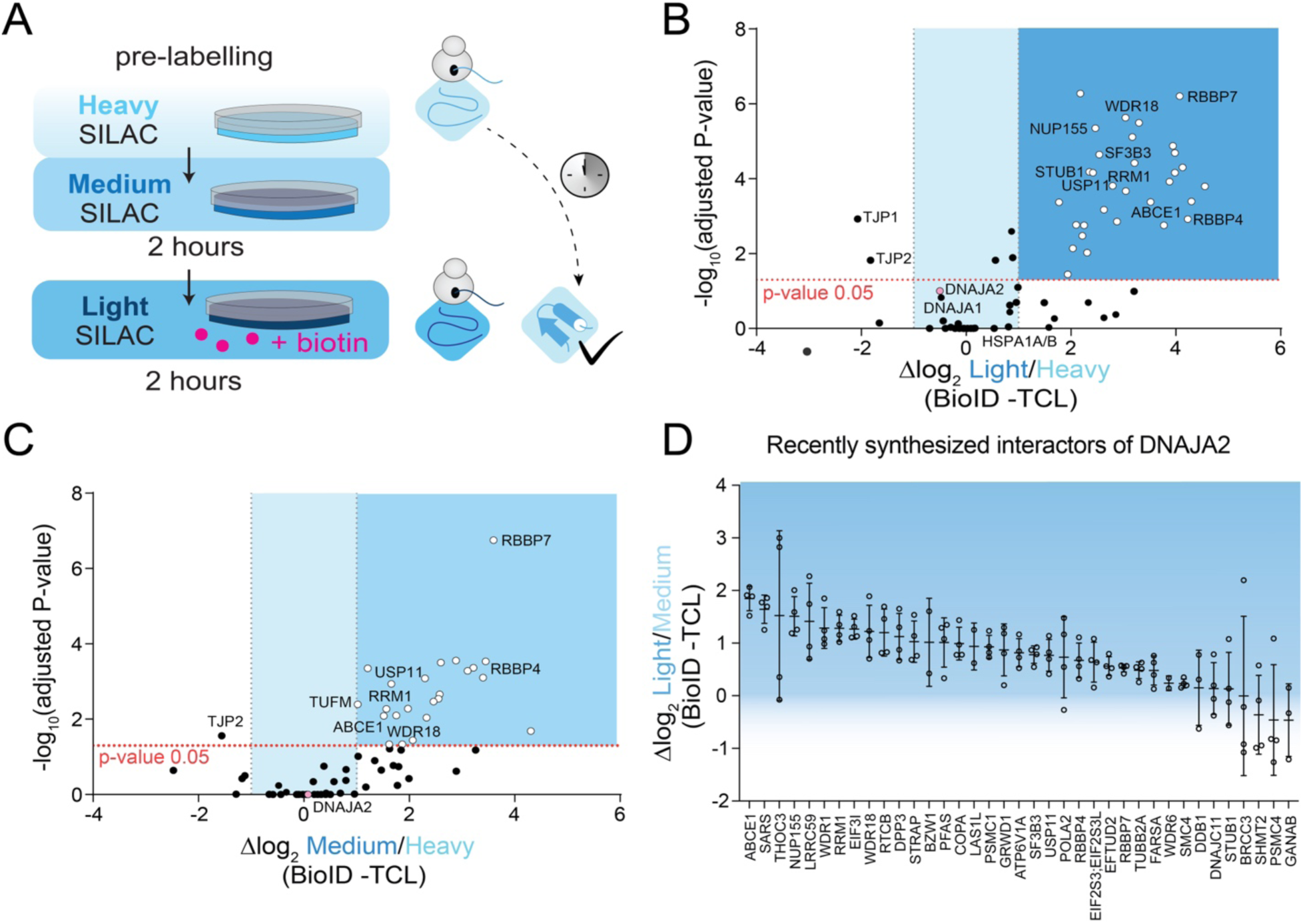
Double pulse-SILAC combined with bioID. **A.** Schematic representation of the double pulse-SILAC labelling workflow in which cells are prelabeled with heavy SILAC (light blue), followed by a pulse with medium SILAC, and then a final pulse with light SILAC (dark blue), during which biotin is also added. **B and C.** Volcano plots showing the log_2_ L/H ratios (B) and the log_2_ M/H ratios (C) of DNAJA2 high-confidence interactors (BFDR<0.01) quantified in the BioID samples and normalized to SILAC ratios in the TCL samples. Statistical significance (p-adj <0.05) was calculated using an unpaired t-test with Holm-Šídák correction. **D.** Plot of the log_2_ L/M values of the recently-synthesized, high-confidence DNAJA2 interactors (BFDR<0.01) quantified in the BioID samples and normalized to SILAC ratios in TCL. Individual values are shown together with mean ratios and standard deviations. Proteins on the left exhibit a stronger enrichment of light versus medium SILAC, whereas proteins on the right represent preys containing a larger fraction of species synthesized during the two-hour period prior to the onset of biotin labelling.

In the TCL, 2331 proteins were quantified in at least two out of four replicates with approximately 15%, 11% and 75% of the protein pools labelled in light, medium and heavy SILAC, respectively (Table S2). Among the 193 DNAJA2 high-confidence interactors identified via SAINT (BFDR <0.01), 67 proteins were quantified in the TCL samples and 37 classified as recently synthesized candidate clients when comparing light-to-heavy (L/H) SILAC ratios (Fig. 2B). When comparing medium-to-heavy (M/H), 34 high confidence interactors remained statistically enriched (Fig. 2C), suggesting the interaction with DNAJA2 may persist over a two-hour period. Nonetheless, when directly comparing light-to-medium (L/M) ratios of the biotinylated proteins, several preys displayed a strong L/M ratio (over twofold) such as NUP155 (Fig. 2D), indicating these candidate clients mostly encountered the co-chaperones within the first two hours of their lifespans. In contrast, the remaining proteins which showed more robust medium SILAC labelling, like as RBBP7, likely remain in close proximity to DNAJA2 for a longer period in time. One advantage of the double pulse labelling is that it also enables clearer separation of recently synthesized preys (light-labelled in this experiment) from pre-existing heavy-labelled proteins.

### DNAJA2 BioID interactors showcase distinct patterns in structural features

Our work thus far reveals that distinct cohorts of proteins engage with the co-chaperone DNAJA2 at different times relative to protein synthesis. We next sought to analyze the features associated with recently synthesized versus other prey proteins. Because only a subset of high-confidence interactors was also quantified in the TCL, we classified preys in the second DNAJA2 experiment based on the overall distribution of ratios in the TCL to avoid bias toward more abundant proteins. Specifically, preys with log_2_ L/H SILAC ratios greater than two standard deviations above the ratios observed in the respective TCL were classified as recently synthesized, whereas all others were not (Fig. 3A). We first observe that both cohorts of DNAJA2 preys are comprised of longer proteins (Fig. 3B) with a lower net charge (Fig. 3C) when compared to proteins quantified in the TCL, as well as the overall proteome. For example, the RBBP7 interactor that we validated has a net charge of −35, which is substantially lower than the proteome-wide average of −2.6. Overall, DNAJA2 candidate interactors do not contain a higher proportion of charged residues, suggesting that the difference in net charge is driven by the distribution and balance of charged residues rather than by their overall abundance. Interestingly, recently synthesized preys are less disordered relative to the TCL and the overall proteome (Fig. 3D). Instead, they contain more β-strands (Fig. 3E) whereas level of α -helices is not different from controls (Fig. S3A). In contrast, pre-existing preys are significantly more disordered (Fig 3D) and exhibit features that are associated with high disorder such as significantly higher PScores (Fig. S3B), which predict the propensity to form pi-pi interactions and are characteristic of proteins that undergo phase separation^45^. These preys have lower melting temperatures (Fig. S3C) and also contain relatively less polar residues (Fig. S3D), a characteristic associated with higher propensity for aggregation and minimizing solvent exposure^46^. Thus, this suggests that these proteins represent polypeptides that continue to engage with the proteostasis network throughout their lifespan to perhaps prevent aggregation.

**Figure 3:**
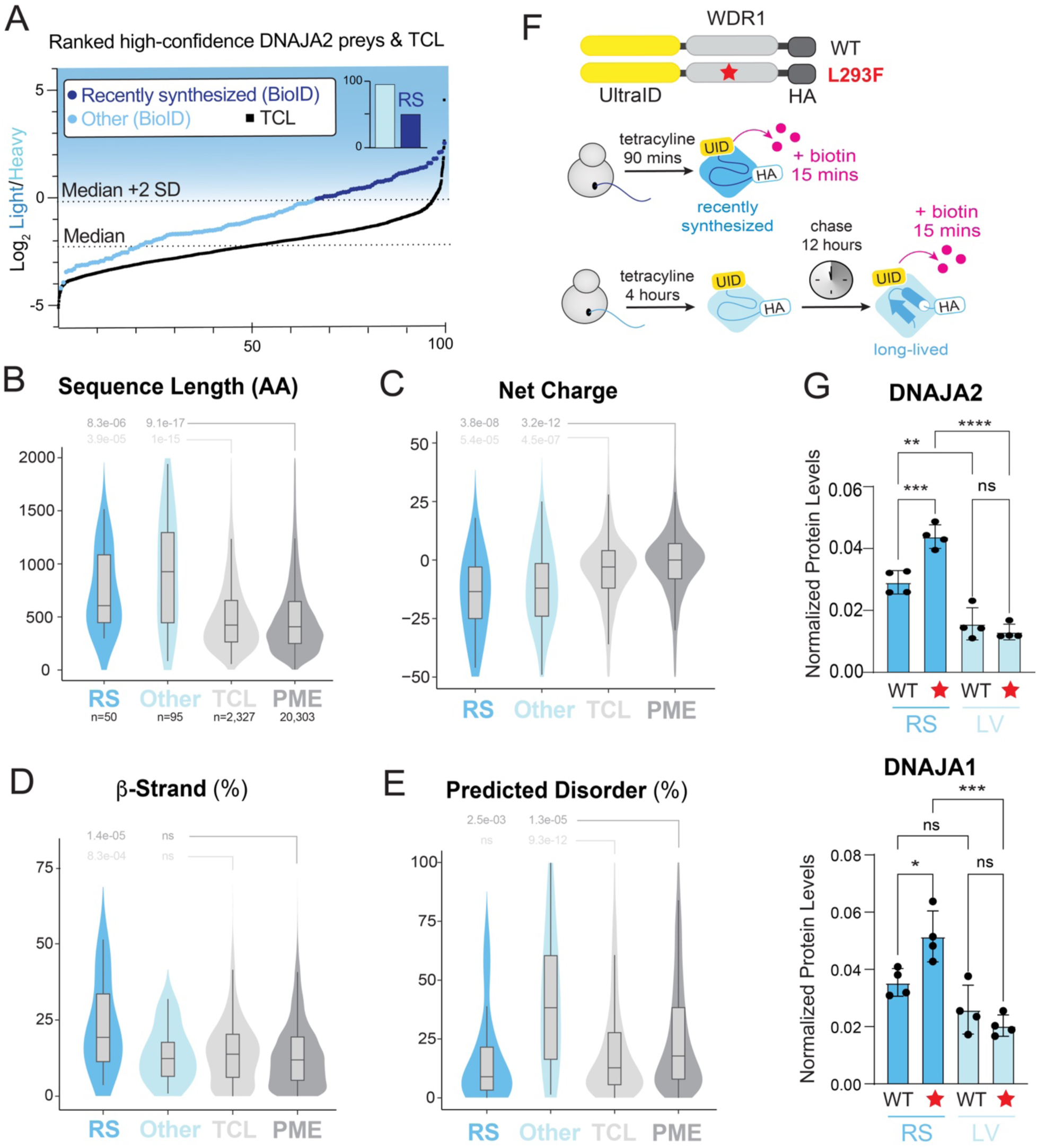
DNAJA2 candidate interactors exhibit distinguished structural features. **A:** Ranked plot of the average log_2_ L/H ratios of proteins identified in the TCL or classified as high-confidence interactors in the BioID dataset. Each datapoint represents a protein quantified in at least two replicates in each condition. Dashed lines indicate the median and +2 standard deviations (SD) of the average log_2_ L/H ratios measured in the TCL. The inset plot shows the distribution of recently synthesized (RS) and all other candidate interactors. **B-E.** Violin plots showing amino acid (AA) sequence length (B), protein net charge (C), percentage of the residues predicted to form β-strand (D) and percentage of residue predicted to be intrinsically disordered (E) among the DNAJA2 high-confidence interactors classified as recently synthesized (RS; n=50) or non-RS (Other, n=95) compared with proteins identified in the TCL or the whole proteome (PME; in D only n=20,301). Statistical significance was assessed using Wilcoxon rank-sum test with Benjamini Hochberg correction for multiple comparisons. **F.** Schematic representation of the reverse-BioID workflow performed in HEK293T cells. UltraID fused to either wildtype (WT) WDR1 or the L293F variant was expressed under a tetracycline-inducible promoter. Following a short 90 min expression period, biotinylation primarily captures proteins interacting with the recently synthesized (RS) WDR1, whereas after the chase period biotinylation predominantly reflects interactions with long-lived (LV) WDR1 species. **G.** Quantifications of DNAJA2 (top) and DNAJA1 (below) levels, normalized to bait levels, following BioID enrichment and DIA analysis using LFQ intensities in the indicated conditions. Statistical analysis was determined by one-way ANOVA with Tukey’s multiple comparison test (GraphPad Prism). Error bars represent mean ± SEM (n=4). **p < 0.01, ***p < 0.001, ****p < 0.0001, ns = not significant.

### Misfolding enhances interaction of recently folded species with DNAJA2

Consistent with the higher levels of β-strands, we noticed that ∼30% of the recently synthesized candidate clients contain WD domains (p-adj <0.05; Table S3). WD domains are characterized by short ∼36-40 amino acid repeats ending with a Trp-Asp dipeptide that assemble into a 7-8 bladed β-propeller structure^47^. These proteins function in various pathways. For example, RBBP7 is involved in histone regulation^48^, whereas WDR1 mediates actin depolymerization^49,50^. Importantly, many of these WD 40 containing interactors are large multi-domain proteins and are established clients of the TRiC chaperonin complex^51^. We reasoned that the biotinylation of recently synthesized proteins captured in our assay could reflect inefficient folding of a subset of these protein pools, thereby requiring multiple rounds of interaction with the proteostasis network well beyond their release from the ribosome.

To test this idea, we employed a reverse-BioID approach to determine whether a misfolded candidate client displays stronger interaction with DNAJA2 when recently synthesized compared to its long-lived counterparts. Specifically, we examined WDR1 carrying the L293F missense mutation in the seventh WD repeat, which is associated with the disease PFIT (period fever, immunodeficiency, and thrombocytopenia)^50^. This mutation disrupts three key hydrogen bonds and impairs closure of the β-propeller structure^50^. The two variants were fused to the UltraID biotin ligase and expressed either for only 90 minutes when proteins start to be detected by Western blots or for 4 hours followed by a 12-hour chase, upon which biotin was added for the last 15 minutes of expression (Fig. 3F). Importantly, levels of the L293F variant were comparable to the wild type (WT) WDR1 in our hands, indicating that the mutation does not substantially affect protein stability (Fig. S4). Cells were then lysed, biotinylated proteins enriched, and samples analyzed by data-independent acquisition (DIA) mass spectrometry for quantitation. Because WDR1 bait levels differed between the recently synthesized and long-lived conditions, prey levels were normalized to their respective WDR1 bait levels to enable direct comparison between conditions. We first observed that recently synthesized WT WDR1 displayed significantly enriched interaction with DNAJA2 relative to its more mature counterpart, confirming our initial findings. Moreover, the L293F variant exhibited significantly increased interaction with DNAJA2 compared to WT WDR1 when recently synthesized, whereas the long-lived pool did not differ (Fig. 3G). Similarly, the recently synthesized L293F variant showed increased interaction with DNAJA1 compared to the recently synthesized WT (Fig. 3G). Together, these results indicate that DNAJA2, perhaps in coordination with DNAJA1, preferentially engages with recently synthesized WDR1 and that misfolding increases this interaction with the co-chaperone. In contrast, mature forms of both variants presumably reach a relatively stable state, leading to reduced interactions with DNAJA2.

### Distinguishing DNAJA2 interactors using pulse SILAC BioID-MS during a heat stress condition

Proteostasis can be disrupted by external stress such as heat shock^52^, and we wondered how chaperone clients may be prioritized in these conditions. We therefore assessed the interactome of DNAJA2 during an acute heat stress at 42°C, a condition in which translation is not quite inhibited^53^. HEK293 cells expressing tetracycline-inducible DNAJA2-FLAG-BioID2 were first pulsed labelled with medium SILAC for two hours followed by heavy SILAC media containing biotin during the heat shock (Fig. 4A). We quantified 2679 proteins in at least two TCL samples with approximately 10%, 10% and 80% of the protein pools labelled with heavy, medium and light SILAC, respectively, consistent with the pattern observed in our previous experiment (Table S4). The presence and distribution of heavy SILAC labelling indicate that translation remained active under these conditions, with no prominent shift toward the synthesis of heat shock-induced proteins in this time frame (Fig S5A, Table S4). Note that we did not further analyze medium-SILAC-labelled proteins, which were included primarily to improve separation between recently synthesized and pre-existing protein species.

**Figure 4:**
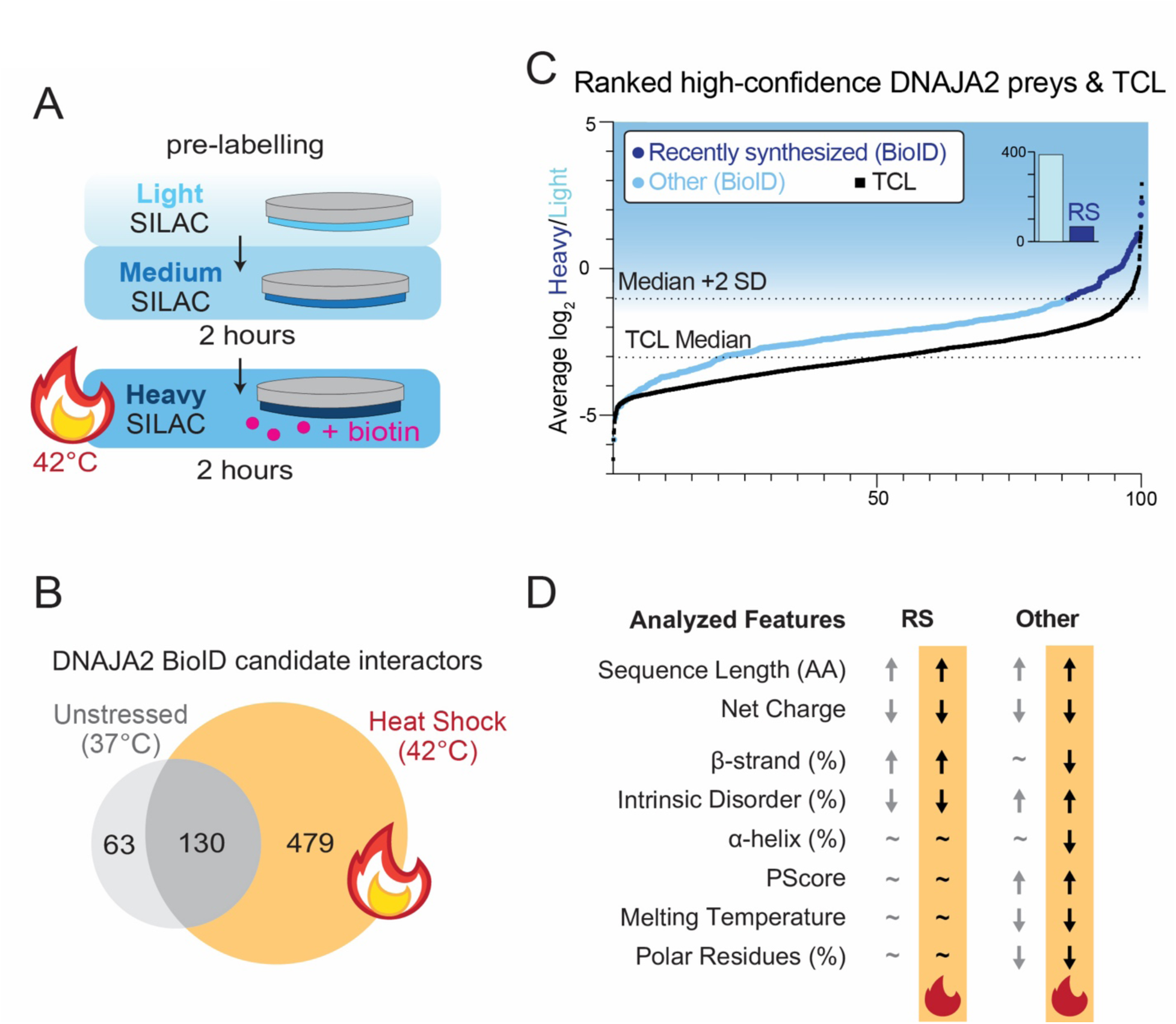
Double pulse-SILAC combined with BioID to resolve interactions under heat shock. **A.** Schematic representing the double pulse-SILAC labelling workflow with heat stress. Cells are prelabeled with light SILAC (light blue), followed by a pulsed with medium SILAC, and then a final pulse with heavy SILAC (dark blue) for two hours in conjunction to biotin treatment, during which the cells subjected to an acute heat stress at 42°C. **B.** Venn diagram of the quantified DNAJA2 high-confidence interactors (BFDR <0.01) in the unstressed (left) and stressed (right) conditions from the two different double pulse-SILAC experiments. **C.** Ranked plot of the average log_2_ H/L ratios of proteins identified in the TCL or classified as high-confidence interactors in the BioID dataset. Each datapoint represents a protein quantified in at least two replicates in each condition. Dashed lines indicate the median and +2 SD of the average log_2_ L/H ratios measured in the TCL. The inset plot shows the distribution of recently synthesized (RS) proteins and all other high-confidence interactors. **D.** Summary of the analyzed features of RS and other DNAJA2 candidate interactors (BFDR <0.01) in the BioID experiments (see also Fig 3, S3 and S5). Arrows indicate a statistically significant enrichment (↑) or depletion (↓) of a given feature relative to the proteome. Non-significant differences are denoted by ∼.

We observed an overall increase in the number of high-confidence DNAJA2 interactors upon the heat shock. A large portion (∼70%) of candidate interactors identified in unstressed conditions were also identified following heat stress (Fig. 4B). Based on the ratio distribution in the TCL, 64 of these candidate interactors were classified as recently synthesized, whereas the remaining 388 preys did not display an increased heavy-SILAC labelling (Fig. 4C). These observations suggest an overall increase in the interaction of misfolded proteins with DNAJA2 during heat stress, with a larger portion of these proteins been synthesized prior to the stress.

We next analyzed the features associated with these two pools of candidate clients, and observed that these properties do not markedly differ from the unstressed conditions (Fig. 4D). Both pools are composed of longer proteins that have lower net charge compared with the TCL and human proteome (Fig. S5B-C). Furthermore, recently synthesized candidate interactors contain significantly more β-strands and are less disordered (Fig. S5D-F). In contrast, the other candidate interactors are significantly more intrinsically disordered and exhibit higher PScores, lower melting temperatures and less polar residues (Fig. S5F-I). Together, these observations indicate that proteins displaying increased biotinylation under heat shock share similar properties with DNAJA2 candidate interactors identified in unstressed conditions, but also that the pool of “older” proteins (i.e., proteins not enriched for heavy SILAC labelling) become more prominent during stress.

## DISCUSSION

We combined pulse-SILAC with BioID mass spectrometry to resolve co-chaperone-client interactions in a temporal manner. More specifically, pulse labelling allowed us to distinguish recently synthesized proteins from pre-existing protein populations that become biotinylated when in close proximity to a co-chaperone. Notably, we found that both these populations are enriched with higher-confidence scoring DNAJA2 interactors that share common characteristics, such as longer sequence length and lower net charge compared to the proteome. In addition, recently synthesized candidate interactors exhibit a higher β-strand content and are frequently enriched in β-propeller domains composed of WD-repeat motifs. In contrast, candidate interactors synthesized prior to pulse metabolic labelling, representing “older” proteins, tend to be more intrinsically disordered and display features associated with a greater propensity for aggregation. Importantly, similar features are retained during a heat stress, although a higher load of pre-existing proteins appear to interact with DNAJA2.

Our study is based on a two-hour metabolic labelling that is required for the detection and quantitation of recently synthesized proteins in the total cell lysate of dividing mammalian cells. However, this labelling window contrasts with the much shorter timescale of co-translational folding, which can occur within minutes. Surprisingly, the double pulse-labelling experiment indicated that the biotinylation of recently synthesized proteins persisted beyond two hours, albeit to a lesser extent. We posit that polypeptides labeled under our conditions may represent a subset of proteins that undergo multiple cycles of association and dissociation with the proteostasis machinery. Interestingly, several WD-repeat-containing proteins enriched in our analysis are established substrates of the TRiC/CCT chaperonin system. Whereas the Hsc70 machinery delivers clients to the TRiC/CCT complex^54^, it is also possible that a subset of proteins that fail to efficiently mature into their native states become increasingly reliant on DNAJA2. Consistent with this notion, the reverse-BioID experiment with the L293F mutant of WDR1 showed a specific increase in association with DNAJA2 following its synthesis compared with the wild type counterpart. Single-molecular tracking approaches, similar to those reported recently^24^ may be required to further elucidate the dynamic trajectories of client protein through successive interaction with the chaperone network.

DNAJA1 and DNAJA2 emerged as high-confidence interactors of one another, consistent with other reports^41,55,56^, and our initial experiments revealed substantial overlap in their recently synthesized interactors. These functional paralogs are well-established co-chaperones of the Hsp70/Hsc70 network across both yeast and mammalian systems, and their overlapping yet distinct functions suggest a coordinated role in the selective recognition of recently synthesized substrates^57,58^. Importantly, when identified as a prey, both proteins displayed metabolic labelling patterns consistent with synthesis preceding the pulse-labelling period. While several other pre-synthesized proteins may be enriched with the two co-chaperones for functional reasons, we posit that a larger fraction of these proteins may instead represent clients. The shift toward a larger enriched pool of these pre-synthesized proteins under heat shock is consistent with this idea. This observation suggests that chaperones may frequently engage proteins long after their synthesis, particularly under conditions of stress. To note, the related cytosolic DNAJA4 was not identified as an interactor for both DNAJA1/A2, either due to its lower levels or its role outside the proteostasis pathways served by DNAJA1/A2.

A key finding is that the inherent features of DNAJA2 interactors differ not only between the recently synthesized and pre-existing proteins, but also from the proteome as a whole, as these proteins are both longer and more negatively charged. One possibility is that DNAJA2 exhibits preferences in client engagement despite its presumed promiscuity and versatility as a co-chaperone. Prior work suggests that DNAJA2 selectively buffers destabilized proteins and recognizes misfolded clients through a beta-hairpin-dependent mechanism^58,59^. In this case, substrate selection may be linked to specific structural features rather than indiscriminate binding. Concurrently or in parallel, a subset of proteins may be more dependent on the proteostasis network. For instance, we previously showed that a subset of proteins that are less disordered and with higher β-strand content compared to the proteome tend to be less thermostable following their synthesis in yeast^25^. These features are reminiscent of the characteristics that distinguish recently synthesized preys from the other subset of high-confidence interactors. Strikingly, similar features were identified among proteins that become more enriched in the Triton-insoluble fraction in brain tissues derived from older mice^60^. This observation suggests that such proteins may not only be particularly dependent on the proteostasis network early in their lifespan but may also impose an increasingly greater burden on the cell as proteostasis capacity declines with age.

Overall, our work showcases how pulse-SILAC coupled with BioID-MS can elucidate the temporal dynamics of chaperone interactions with recently synthesized and pre-existing proteins. Better understanding of chaperone function will help clarify how proteostasis is maintained and how protein quality control mechanisms contribute to the prevention of disease.

## ACKNOWLEDGEMENTS

We are grateful for Dr. Mikko Taipale for sharing with us the pcDNA5 FRT/TO Gateway destination and pDONR223 Gateway entry vectors and reagents, as well as for the insightful discussions. We are grateful for Jason Rogalski and the UBC Proteomics Core Facility for assistance with the proteomics experiments, and Jenny Moon from the UBC Foster lab for supporting us with the SILAC reagents used in this study. We also thank Dr. Anne Claude Gingras for gifting us the psTV6-N-UltraID plasmid. Lastly, we also thank past and present members of the Mayor lab for productive discussions that drove the work done in this study, including Dr. Molzahn for exploratory work not included in this manuscript and Dr. Kuechler for sharing precalculated feature data.

## Author Contributions

Experimental Design and Methodology: S.K, G.C and T.M; Experimental work: S.K, G.C, V. E.P, C.S.D, H.E and E.H; Data analyses: S.K, G.C and T.M; Resources: S.C, G.C; Supervision: T.M; Writing (first drafts): S.K and T.M; Editing: G.C, S.K and T.M.

## METHODS

### Molecular Cloning and Plasmids Used in This Study

All plasmids used and generated in this study are outlined in Table S5. Gateway LR II Clonase Mix (Invitrogen) was utilized with manufacturer’s instructions to generate the pcDNA5 FRT/TO HSPA1A-FLAG-BioID2, pcDNA5 FRT/TO DNAJA1-FLAG-BioID2, pcDNA5 FRT/TO DNAJA2-FLAG-BioID2 and pcDNA5 FRT/TO RBBP7– 3x HA constructs. The pcDNA5 FRT/TO ccdB Gateway Destination and pDONR223 Entry clones used for the above cloning were constructs provided kindly by Dr. Mikko Taipale. The psTV6-N-UltraID construct, was a gift from Dr. Anne Claude Gingras which was used to generate the pcDNA5 FRT/TO UltraID-3xHA-WDR1 and UltraID-3xHA construct using Gibson Assembly (New England Biolabs). pcDNA5 FRT/TO UltraID-3xHA-WDR1-L239F mutant was generated using site directed mutagenesis (New England Biolabs Base Changer tool) using a kinase, ligase and Dpn1 reaction mix. Whole plasmid sequencing (Plasmidsaurus) was utilized to verify generation of the correct plasmid vectors.

### Cell culture for generation of Stable Cell Lines

Flp-In™ T-Rex™ HEK 293 cells were cultured in Dulbecco’s Modified Eagle Medium (DMEM) (Gibco) supplemented with 1% penicillin-streptomycin (Gibco) and 10% fetal bovine serum (FBS) (Gibco) at 37°C and 5% CO_2_. Routine passaging was carried out using 0.25% Trypsin-EDTA (Gibco) and cell counting was performed using the automated cell countess II instrument (Thermofisher Scientific). To generate stable Flp-In™ T-Rex™ HEK 293 cells expressing tetracycline-inducible fusion proteins, cells were first grown to 75-80% confluency in 6 cm tissue culture dishes. Cells were then then co-transfected with 2µg of pOG44 Flp recombinase expression Vector (Invitrogen) and 1µg of the respective pcDNA5 FRT/TO construct using FuGENE 6 (Promega) at a 3:1 FuGENE:total plasmid ratio. 24 hours following transfection, cells were expanded to a 10 cm dish and further grown for 24 hours, upon which Hygromycin B (5 μM; BioShop) was added. Continued selection with Hygromycin B was carried out every 3-4 days until visible colonies (∼2-4mm diameter) of polyclonal populations covered the dish. Then, colonies formed on one half of the plate were harvested and transferred to a new 10 cm dish (population A), while colonies from the other half were harvested and transferred into a separate 10 cm dish (population B). Each clone population generated was expanded to generate more replicates for downstream experiments.

### Cell culture for BioID experiments

For all BioID experiments, two replicates of each population A and population B were subjected to the workflow to generate four replicates.

#### For single pulse-SILAC BioID-MS (Experiment 1)

Stable Flp-In™ T-Rex™ HEK 293 cells expressing tetracycline-inducible protein of interest (POI)-FLAG-BioID2 proteins were grown in light SILAC media (DMEM, 10% dialyzed FBS (Gibco), 1% penicillin-streptomycin (Gibco), 45.62mg/L Lys-0, 36.88mg/L Arg-0) and seeded at an approximate density of 1.6 × 10^6^ cells unto 10 cm dishes. When cells were ∼50% confluent, tetracycline (1μg/mL) was added, and cells were further incubated for 24 hours. Cell culture media was then switched to heavy SILAC media (DMEM, 10% dialyzed FBS (Gibco), 1% penicillin-streptomycin (Gibco), 45.62mg/L Lys-8 (Silantes), 36.88mg/L Arg-10 (Silantes) containing biotin (50μM) for 2 hours. Cells were then harvested by scraping into 1xPBS, pelleted (2 min, 1000 r.c.f, 4°C), snap frozen with lN_2_ and stored at −70°C until later use.

#### For double-pulse SILAC BioID-MS (Experiment 2)

Stable Flp-In™ T-Rex™ HEK 293 cells expressing tetracycline-inducible POI-FLAG-BioID2 proteins were grown in heavy SILAC media in 35 mm dishes for at least 6 doublings to ensure proper labelling and seeded at an approximate density of 1.6 × 10^6^ cells unto 15 cm dishes. When cells were ∼50% confluent, tetracycline (1μg/mL) was added, and cells were further incubated for 24 hours. Cell culture media was then switched to medium SILAC media (DMEM, 10% dialyzed FBS (Gibco), 1% penicillin-streptomycin (Gibco), 45.62mg/L Lys-4, 36.88mg/L Arg-6) for 2 hours. Then, cell culture media was switched to light SILAC media containing biotin (50μM) for 2 hours. Scraped cells were pelleted (3 min, 1000 r.c.f, 4°C), washed twice with 1x PBS, snap frozen with lN_2_ and stored at −70°C until later use.

#### For double-pulse SILAC BioID-MS with heat shock (Experiment 3)

Stable Flp-In™ T-Rex™ HEK 293 cells expressing tetracycline-inducible POI-FLAG-BioID2 proteins were grown in light SILAC media and seeded at an approximate density of 1.6 × 10^6^ cells unto 15 cm dishes. When cells were ∼50% confluent, tetracycline (1μg/mL) was added, and cells were further incubated for 24 hours. Cell culture media was then switched to medium SILAC for 2 hours. Then, cell culture media was switched to heavy SILAC media containing biotin (50μM) and cells were subjected to a heat shock at 42°C for 2 hours. Heat shock was carried out by incubating cell culture dishes in a pre-heated water bath, inside a 42°C cell culture incubator. Cells were harvested as done in Experiment 2.

#### For reverse pulse-chase BioID

Stable Flp-In™ T-Rex™ HEK 293 cells expressing tetracycline-inducible UltraID-3xHA-WDR1 (WT and L293F mutants) were seeded at 1.2 × 10^6^ cells in 10cm dishes. At ∼70% confluency, tetracycline (1μg/mL) was added for 4 hours to induce protein expression of long-lived UltraID-3xHA-WDR1, and then tetracycline was removed by changing the media and further incubation (chase) for 12 hours before biotin treatment. For expression of recently synthesized UltraID-3xHA-WDR1, tetracycline (1μg/mL) was added at ∼90% confluency for 90 minutes. Biotin treatment (50μM) was done in the last 15 minutes of the 90-minute tetracycline treatment or of 12-hour chase, for the recently synthesized condition and long-lived condition, respectively. Cells were then collected via trypsinization and centrifugation (2 min, 1000 r.c.f, 4°C) and then cell pellets were washed twice with 1xPBS, snap frozen with lN_2_ and stored at −70°C until later use.

### Reverse Pulse-Chase Co-Immunoprecipitation

Stable Flp-In™ T-Rex™ HEK 293 cells expressing tetracycline-inducible RBBP7-3xHA were seeded at 1.2 × 10^6^ cells in 10 cm dishes. At ∼70% confluency, tetracycline (1μg/mL) was added for 6 hours to induce protein expression of long-lived RBBP7-3xHA, and then tetracycline was removed by changing the media and further incubation (chase) for 18 hours before biotin treatment. For expression of recently synthesized RBBP7-3xHA, tetracycline (1μg/mL) was added at ∼90% confluency for 2 hours. Cells were then collected via trypsinization and centrifugation (2 minutes, 1000 r.c.f, 4°C) and then cell pellets were washed twice with 1xPBS and then kept on ice until use. Protein crosslinking was done by adding 1mL of 2mM DSP (dithiobis(succinimidyl propionate)) (2mM in 1xPBS; Thermo Fisher Scientific) to cell pellets, and incubated end-over-end rotation (40 minutes, RT). The reaction was quenched with 100µL of Tris-HCl (50mM final concentration, pH 7.4) with continued mixing (15 minutes, RT). Cells were pelleted via centrifugation (2 minutes, 1000 r.c.f, 4°C), washed once with 1x PBS, and kept on ice until use. Cell lysis was carried out with 1mL of HEPES lysis buffer (50 mM HEPES, 150 mM NaCl, 1% NP-40, and freshly added 1mM PMSF and 1xPIC) and passaged with a 27G needle 5 times and incubated on ice for 20 minutes. Cell debris was cleared by centrifugation (15 minutes, 15,000 r.c.f., 4°C), and soluble supernatant was collected for BCA assay. Lysates were brought to 1 μg/μL in a total of 1000 μL, and 38 μg was saved aside as the input fraction. The rest of the lysate was incubated with TBS-T equilibrated Pierce™ anti-HA Magnetic Beads, and the co-immunoprecipitation was carried out by closely following the manufacturer’s protocol^61^ with minor modifications: washes were carried out with TBS-T containing 1M NaCl, and elution was done with 60μL of HA-peptide (Thermo Fisher Scientific).

### Western Blots

Proteins were diluted with Laemmli buffer and separated on either 4-20% Mini-PROTEAN® TGX™ Precast Protein Gels (Bio-Rad) or home-cast 10% SDS-PAGE gels. Semi dry transfer was done via 0.45μm nitrocellulose membrane with the TransBlot® Turbo^TM^ Transfer System (Bio-Rad) standard protocol. Membranes were blocked with 5% powder milk in TBS-T, and incubated with primary antibody (diluted in 5% milk in TBS-T) overnight at 4°C. 1x TBS-T was used for washes, and secondary antibody (diluted in 5% powder milk in TBS-T) incubation was carried out for 30 minutes at room temperature. Imaging and quantitation of blots was done using the Odyssey (Li-Cor). The following primary antibodies were used: anti-biotin (AB53494, Abcam, 1:1,000), anti-GAPDH (14C10, Cell Signaling Technology; 1:5,000), anti-HA (51064-2-AP, Proteintech, 1:1000), anti-DNAJA2 (12236-1, Proteintech), anti-alpha-tubulin (66031-1, Proteintech, 1:1000). The following secondary antibodies were used: IRDye 800CW Goat anti-Mouse (Li-Cor; 1:10,000), IRDye 680RD anti-rabbit (Li-Cor; 1:10,000) and IRDye 800CW Streptavidin (Li-Cor; 1:10,000).

### Immunofluorescence

Flp-In™ T-Rex™ HEK 293 cells expressing tetracycline-inducible POI-FLAG-BioID2 were seeded at 0.2 × 10^6^ cells in poly-D-lysine coated coverslips in 6-well plate, with media containing tetracycline (1μg/mL). After 24 hours, biotin (50μM) was added for another 24 hours. Cells were then fixed with 1mL of 4% Formaldehyde (w/v) Methanol-free (Pierce) (10 minutes, room temperature). Fixed cells were washed thrice with 1xPBS, and then permeabilized using 2mL of 0.5% Triton-X in 1x PBS (2 minutes, room temperature). Washes were repeated three times with 1x PBS and blocked with 3% BSA in 1xPBS (1 hr, room temperature). Incubation with primary antibodies and Hoescht 33342 (Invitrogen, 1:10,000 in 1xPBS) was carried out overnight at 4°C. Washes were repeated thrice with 1x PBS and then incubated with fluorescently labelled secondary antibodies (45 minutes, room temperature). Washes with 1xPBS were repeated thrice and coverslips were mounted on slides using ProLong™ Gold Antifade Mountant (Invitrogen) and sealed using nail polish. Images were obtained on an AXIO Observer 7 inverted led fluorescence microscope (Zeiss) and processed with the Zen Blue 3.0 image processing software. The primary antibodies used were anti-biotin (AB53494, AbCam; 1:1,000) and anti-FLAG (F1804-50UG, Sigma Aldrich, 1:1,000). The secondary antibodies were Alexa Fluor^TM^ 594 (anti-mouse) and anti-Alexa Fluor^TM^ 488 (anti-rabbit) (Invitrogen, 1:10,000).

### Streptavidin-Affinity Purification of Biotinylated Proteins and Sample prep for MS

#### Pulse-SILAC-BioID

Cell pellets were first thawed on ice and lysed with 1mL (for experiment 1) or 1.8mL (for experiment 2 and 3) of Triton lysis buffer (50mM Tris pH 7.5, 150mM NaCl, 1% Triton) freshly supplemented with cOmplete, EDTA-free Protease Inhibitor Cocktail (1x, Roche) and PMSF (1μM). Lysates were incubated on ice for 30 minutes and then cleared for cell debris by spinning (for experiment 1: 15 minutes, 900 r.c.f, 4℃; for experiments 2 and 3: 20 minutes, 16,000 r.c.f, 4℃). The soluble supernatants were transferred to a fresh lo-bind tubes (Sarstedt) and protein concentrations were measured with a BCA assay (Thermofisher) according to the manufacturer’s protocol without modifications. 100μg of protein lysate was kept aside for total protein analysis carried out by SP3 digestion as explained below. All lysates were normalized to the same concentration and volume, then each was added to 30 µL of High-Capacity Streptavidin Agarose (Pierce) beads pre-equilibrated in 50 µL Triton lysis buffer and then incubated overnight at 4°C on an end-over-end rotator. Beads were then loaded to spin columns (Pierce™ Spin Columns - Snap Cap #69725) and washed twice with triton lysis buffer via centrifugation (750μL, 500 r.c.f, 4℃). The following steps were done at room temperature unless indicated otherwise. Beads were washed twice with 0.5x SDS in 1xPBS (750μL, 500 r.c.f) via centrifugation, and then incubated with 1xPBS SDS solution containing 100mM DTT (20 minutes, 100μL for experiment 1, 200μL for experiments 2 and 3). Next, beads were washed ten times with UC buffer (750μL; 6M urea, 100 mM Tris-HCl pH 8.5), and then incubated with UC buffer containing 50mM iodoacetamide (20 minutes; 200µL) in the dark. Following that, samples were washed ten more times with UC buffer as done prior and then thrice with water (750µL). Beads were then resuspended in ammonium bicarbonate (50mM) containing 1µg of sequencing grade trypsin (Promega)(100µL), and spin columns were sealed with parafilm. Trypsin digestion was carried out at 37°C overnight and peptides were collected into fresh lo-bind tubes via centrifugation (1min, 1000 r.c.f).

For total cell lysates, Single-pot, solid-phase-enhanced sample preparation (SP3) was carried out^62^. 100µg of TCL was reduced with Tris(2-carboxyethyl)phosphine (TCEP) (3mM) at 25°C for 20 minutes with shaking at 900 rpm, then alkylated with 2-chloroacetamide (55mM) at 25°C for 20 minutes with shaking at 900 rpm. Ethanol (100%) was to the samples at a 1:1 ratio, then magnetic beads (50% SeraMag A and 50% SeraMag B) were added a ratio of 1:10 mg protein: mg beads and incubated at 25°C for 10 min, shaking at 900 rpm. Next, beads were isolated by a magnetic plate, washed thrice with 80% Ethanol and resuspended in ammonium bicarbonate (50mM) containing trypsin at a 1:50 ratio, incubated at 37°C overnight with shaking at 1000 rpm.

Peptides were acidified with 10% TFA prior to C18 clean up and MS analysis. Peptide desalting was performed manually, closely following the protocol by Rappsilber et al^63^. Briefly, 6µg of high-capacity C18 (Phenomenex) StAGE Tips was rehydrated sequentially using 100% Methanol, Buffer B (80% acetonitrile, 0.1% TFA), and Buffer A (5% acetonitrile, 0.1% TFA), centrifuging at 1000 rcf for 5 min at room temperature at each step while keeping the resin hydrated. Acidified peptides were loaded and columns were washed with Buffer A then stored at 4 °C if needed. Peptides were eluted with Buffer B2 (40% acetonitrile, 0.1% TFA), vacuum-dried, and stored at −20 °C until injection. Dried peptides were reconstituted in MS buffer (0.1% formic acid, 0.5% acetonitrile) prior to injection.

#### Reverse Pulse-Chase BioID

Cell pellets were first thawed on ice and lysed with 300μL of modified RIPA buffer (50mM Tris pH 7.5, 150mM NaCl, 1% Triton, 0.5% Sodium Deoxycholate, 0.1% SDS) freshly supplemented with cOmplete, EDTA-free Protease Inhibitor Cocktail (1x, Roche) and PMSF (1μM). Lysates were incubated on ice for 25 minutes and then sonicated using a water bath sonicated (Bioruptor) for 5 minutes (pulses of 30s on, 30s off on High). Cell lysates were cleared of cell debris by centrifugation (15 minutes, 15,000 r.c.f, 4℃) and the soluble supernatant was collected in fresh lo-bind tubes. Following a BCA assay, samples were normalized to 5μg/μL for BioID pulldown. The entirety of the pulldown was carried out on the BRAVO liquid handler with 96LT head (Agilent) using an in-house protocol modified from Lam et al^64^and developed in-house on the VWorks software for automation. Briefly, for each sample, 600μg of lysate was enriched with 6μL of Streptavidin Magnetic Beads (Pierce) pre-equilibrated in modified RIPA lysis buffer (90 minutes, 4℃). Beads were then washed twice with modified RIPA lysis buffer (100μL), once with KCl (1M; 100μL), twice again with modified RIPA lysis buffer (100μL) and once with 1xPBS(100μL). All washes were done at room temperature with shaking incubations (10 minutes, 1600 rpm) for each wash. Post-washes, the beads were resuspended with 50mM ammonium bicarbonate and supplemented with DTT (10mM final concentration) for 30 minutes, then with 2-chloroacetamide (55mM final concentration) for another 30 minutes. To this mixture, on-bead trypsin digestion (overnight on Thermomixer R (Eppendorf), 1400 rpm, 37℃) was done by adding a total of 0.2μg of sequencing-grade trypsin per sample. Peptides were collected by separating the beads using a magnetic 96-well plate (Permagen Labware) and were acidified with 10% TFA. Peptide desalting (C18 clean up) was then done via the StAGE-Tipping protocol automated on the BRAVO liquid handler (Agilent) using the Peptide cleanup utility with AssayMap head and C18 cartridge tips (Agilent). Desalted peptides were dried using a SpeedVac (Eppendorf) concentrator and stored at −20°C. Dried peptides were reconstituted in MS buffer.

### Mass spectrometry

#### Pulse-SILAC-BioID

Each biological replicate was injected once, except for the RLuc samples, where each biological replicate was loaded 4 times. For each, 120ng of guesstimated peptides were injected and separated on-line using Easy-nLC 1200 (Thermo Fisher Scientific) with Aurora Series analytical column, (25cm x 75μm 1.6μm C18; Ion Opticks, Parkville, Victoria, Australia). The analytical column was heated to 40°C using an integrated column oven (PRSO-V2, Sonation, Biberach, Germany). Buffer A consisted of 0.1% aqueous formic acid and 2% acetonitrile in water, and buffer B consisted of 0.1% aqueous formic acid and 80% acetonitrile in water. A standard 90 min gradient was run from 2% to 25% buffer B over 46 min, then to 35%, 50% and 95% buffer B over 15, 5 and 5 min, respectively, to be held at 95% buffer B for 8 min before dropping to 3% over 2 min to be for another 6 min. The analysis was performed at 0.25 μL/min flow rate. The Easy-nLC thermostat temperature was set at 7°C. The peptides were analyzed with an Orbitrap Exploris 480 mass spectrometer (Orbitrap Exploris^TM^ 480, Thermo Fisher Scientific). The Nanospray Flex^TM^ ion source was operated at 1900 V spray voltage and ion transfer tube was heated to 290°C. During analysis, the Orbitrap Exploris 480 was operated in a data-dependent acquisition (DDA) mode. The parent scan MS resolution was set to 120000, with 100% normalized automatic gain (AGC) target, 50% RF lens, 20ms maximum injection time and scan range from m/z 375Th to m/z 1200Th. Then, the top 20 most abundant precursor ions were selected for fragmentation by a higher-energy collisional dissociation (HCD) normalized collision energy of 28%. MS/MS spectra were collected at a resolution of 15000, normalized AGC target of 50%, and an isolation window of m/z 2Th. Dynamic exclusion was set so that after a single MS/MS detection, precursor ions within a 10ppm mass tolerance were excluded from selection for 25s. Protein identification was carried out by MaxQuant^65^(v2.1.0.0) using the UniProt reviewed human FASTA file supplemented with, streptavidin, BioID2, and Renilla luciferase sequences. Search parameters were set to default; variable modifications for Oxidation [+16] on M and Acetyl [+42] at protein N-terminus and maximum five variable modifications per peptide were allowed. The fixed modification was Carbamidomethyl [+57] on C. Trypsin/P was used as the specific enzyme, peptide length selected as 7 to 52 with a maximum of two missed cleavages. Standard multiplicity was selected, and Lys8, Lys4, Lys0, Arg10, Arg6 and Arg0 were selected for identification of SILAC labelling.

#### Reverse Pulse-Chase BioID

75ng of guestimated peptides were injected in a similar manner for analysis with the Orbitrap Exploris^TM^ 480 operated in a DIA mode. A standard 90 min gradient was run from 2% to 20% buffer B over 46 min, then to 32%, 50% and 95% buffer B over 15, 5 and 5 min, respectively, to be held at 95% buffer B for 8 min before dropping to 3% over 2 min to be for another 6 min. Full scans MS resolution was set to 60,000 with 300% normalized automatic gain control (AGC) target, 50% RF lens, 25 ms maximum injection time and scan range from m/z 380Th to m/z 985Th. DIA fragment spectra were collected at a resolution of 15,000, normalized AGC target of 2000%, maximum injection time of 40 ms, scan range from m/z 145Th – m/z 1450Th. Isolation windows of m/z 10Th were used with an overlap of m/z 1 Th. Normalized collision energy was set to 28%. The mass-to-charge ratio was calibrated based on three selected ions from Pierce^TM^ FlexMix^TM^ Calibration Solution (m/z [Th]: 195, 322, 524, 622, 922, 1122, 1222, 1322, 1422, 1522, 1622, 1722 and 1822). The mass accuracy was typically within 2ppm and was not allowed to exceed 4ppm. Protein identification was carried out by Spectronaut direct DIA pipeline (v19.8.250311.62635; Biognosys) using a UniProt reviewed human FASTA file supplemented with streptavidin (UniprotID – P22629), trypsin (UniprotID – P00761) and UltraID sequences. Search parameters were set to default. Protein identification for SAINTq analysis was done via DIA-NN ^66^(v2.0.2) using an in silico spectral library generated with the same FASTA file as above. The raw fragment quantities were exported by adding the “-- export-quant” command in the additional options box.

### Data analysis and Statistical Tests

#### Pulse-SILAC-BioID

Significance Analysis of INTeractome (Saintq v0.0.4^67^) was used to identify high-confidence interactors. Intensities of quantified proteins for each labelling channel were summed, and reported sum of intensities of each channel was used for the SAINT, using the top four intensities from the RLuc replicates as the background controls. Both test and control baits used a replicate compression setting of *compress_n_rep = 100.* Cut-offs of BFDR< 0.05 in the pilot and < 0.01 in the double SILAC experiments were selected (a more stringent cutoff was applied in the double SILAC experiments due to the higher number of candidate preys). SILAC ratios of all identified proteins were log_2_ transformed in Excel. To compare BioID with TCL samples, GraphPad Prism (v11.0.1) was used for assessing statistical significance using a multiple unpaired t-tests with Holm-Šídák method for multiple hypothesis corrections, as well as to generate volcano plots. Only interactors that were quantified in at least two of the four biological replicates were used for feature analysis and statistical analysis which was done using an in-house generated R-script for analysis in previous work ^68,69^. Briefly, the feature analysis was done for the sequence length, net charge and percent polar residues obtained with Perl (v5.22.1)^70^, intrinsic protein disorder calculated with DISOPRED3^71^, protein melting temperatures measured in Leuenberger et al^72^. PScores calculated in Vernon et al^73^, secondary structure (β-strands and α-helices content) calculated by SSpro in the SCRATCH-1D 1.1 software package^74^and with PSIPRED^75^.The nonparametric Mann–Whitney U test with Benjamini Hochberg’s multiple hypothesis correction to identify significance of features in high-confidence interactors relative to the total human proteome (p-adjusted value <0.05).

#### Reverse Pulse-Chase BioID

PG.Quantity values from the Spectronaut report served as the protein intensities for assessing chaperone levels in each sample condition. Levels of DNAJA2 were normalized to levels of WDR1 in the respective sample, and GraphPad prism (v11.0.1) was used to perform a two-way ANOVA test with Tukey’s correction between levels in WDR1 WT vs L293F mutant, in recently synthesized and long-lived conditions.

### Resources

- Raw MS files are deposited in the MassIVE repository (ftp://MSV000102243@massive-ftp.ucsd.edu; doi:10.25345/C5G15TR59
- All code used in the analysis are deposited at - https://github.com/shriyakamat/pulseSILAC_BioID_Feature_Analysis_Pipeline.
- Any data associated with the studies are available from the corresponding author upon request.

---

**Figure S1:**
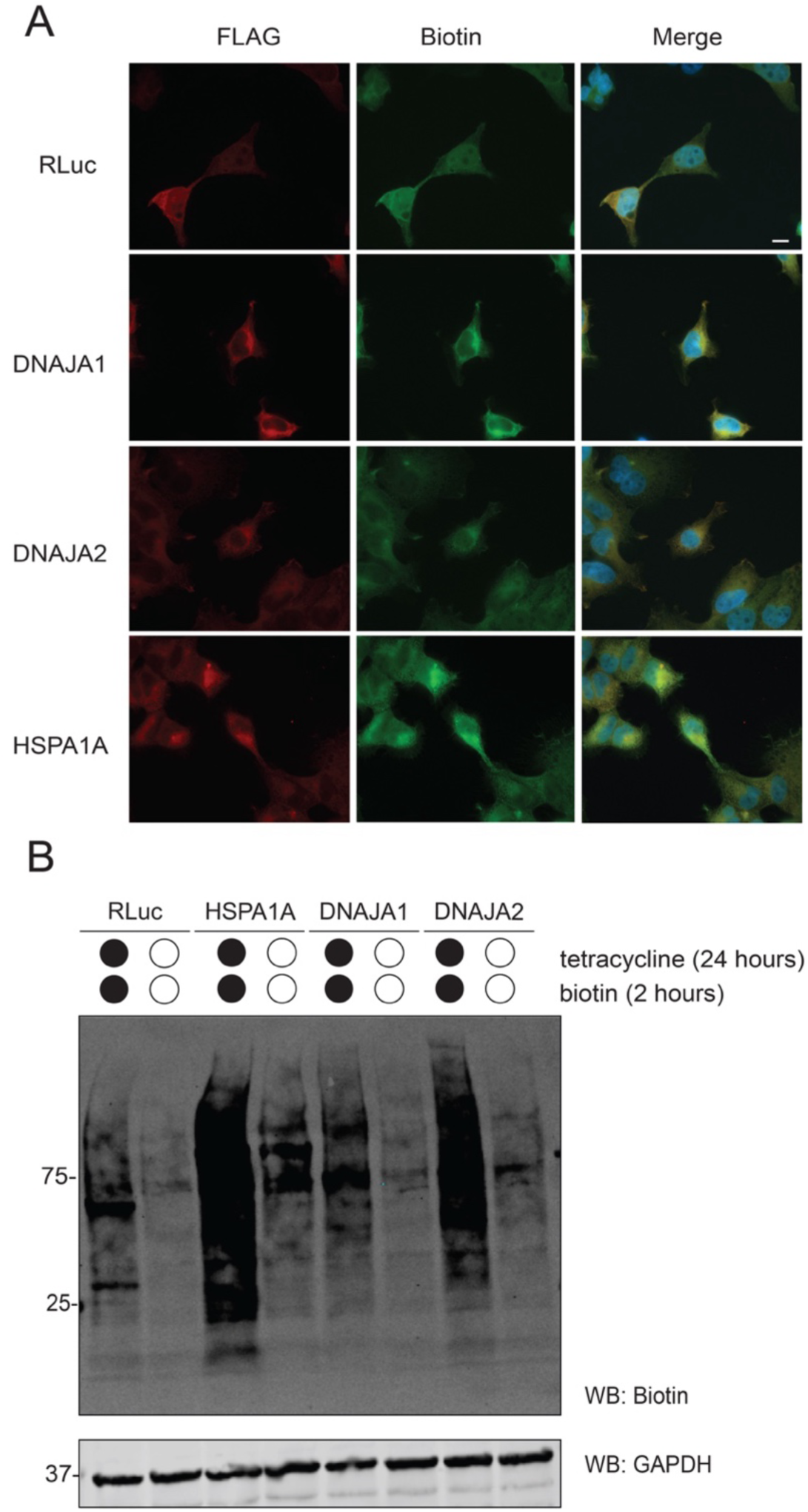
Tetracycline-inducible BioID2 constructs show sufficient biotinylation signal at two hours and appropriate cytosolic localisation. **A.** Immunofluorescence images showing the subcellular localization of the BioID2-fused control, RLuc, and the co-chaperones DNAJA1 and DNAJA2 in HEK293 cells, detected using anti-FLAG staining, together with the corresponding biotinylation patterns detected using anti-biotin staining. Nuclear staining is indicated with the Hoechst stain. Scale bars: 10μm. **B.** Western blot analyses of short biotinylation treatment. Expression of BioID2 fused baits were induced via tetracycline treatment (1μg/mL) for 24 hours, followed by a biotin treatment (50μM) for 2 hours (black circles). Untreated samples (empty circles) serve as control lanes. GAPDH is shown as the loading control.

**Figure S2:**
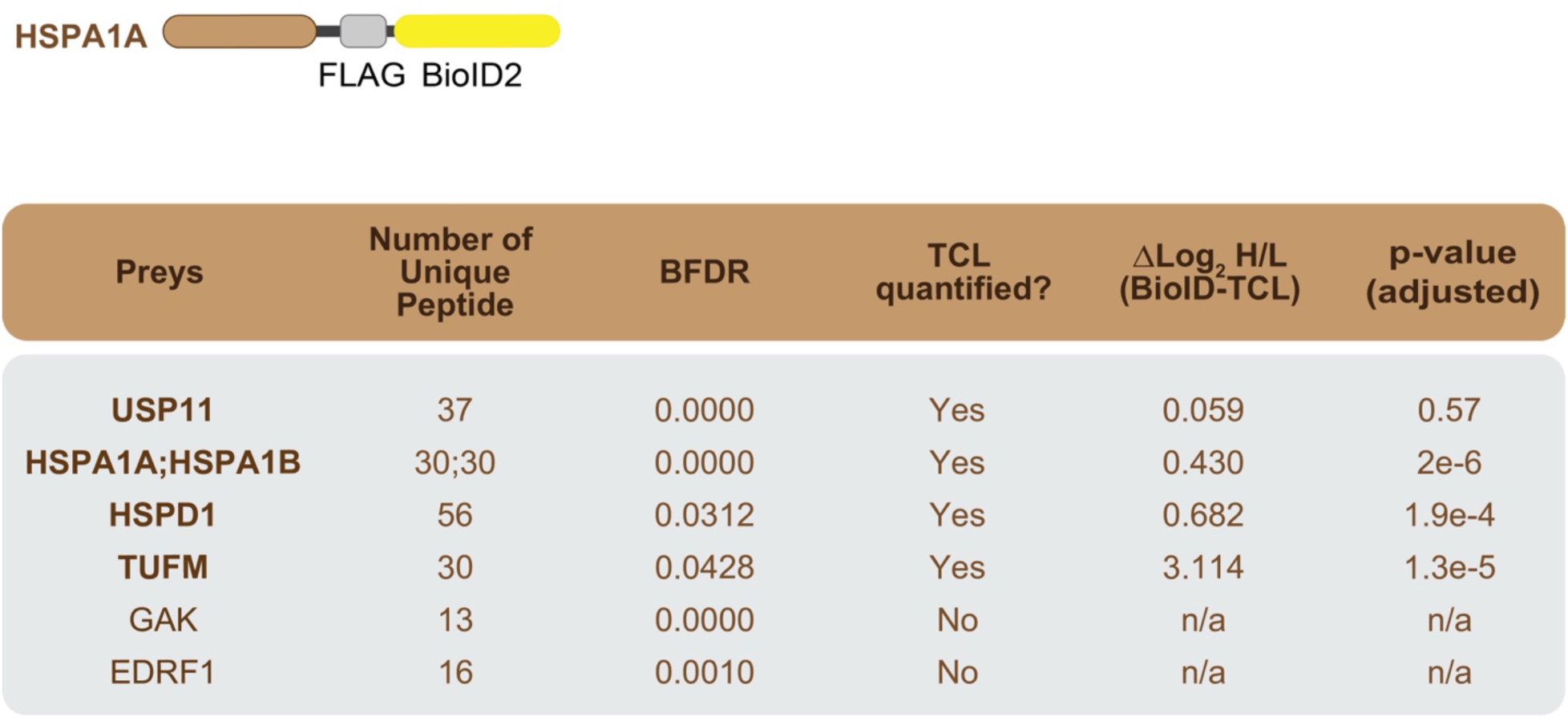
Single pulse-SILAC BioID-MS experiment with HSPA1A. Schematic of the HSPA1A-FLAG-BioID2 fusion protein and list of high-confidence HSPA1A interactors identified by BioID (BFDR <0.05), together with the number of unique peptides detected, SAINTq BFDR values, and TCL-normalized log_2_ SILAC ratio if applicable.

**Figure S3:**
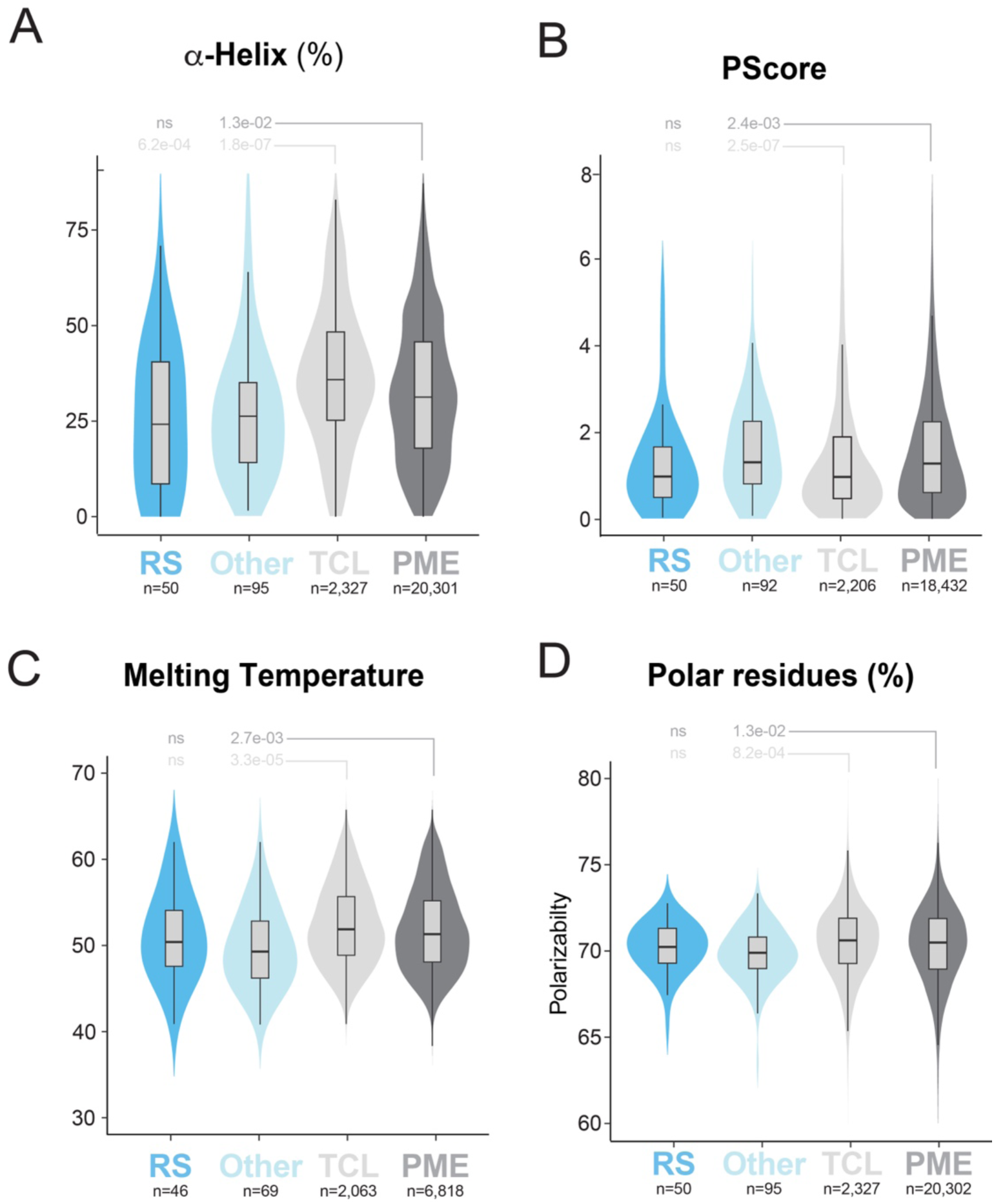
Recently-synthesized and other DNAJA2 high-confidence interactors exhibit distinct structural features. **A–D.** Violin plots showing the percentage of residue predicted to form α-helices (A), protein melting temperatures (B), PScores for pi-pi interactions (C), and the percent polar residues (D) among the DNAJA2 high-confidence interactors classified as recently synthesized (RS; n=50) or non-RS (Other, n=95) compared with proteins identified in the TCL or the whole proteome (PME). Statistical significance was assessed using Wilcoxon rank-sum test with Benjamini Hochberg correction for multiple comparisons.

**Figure S4:**
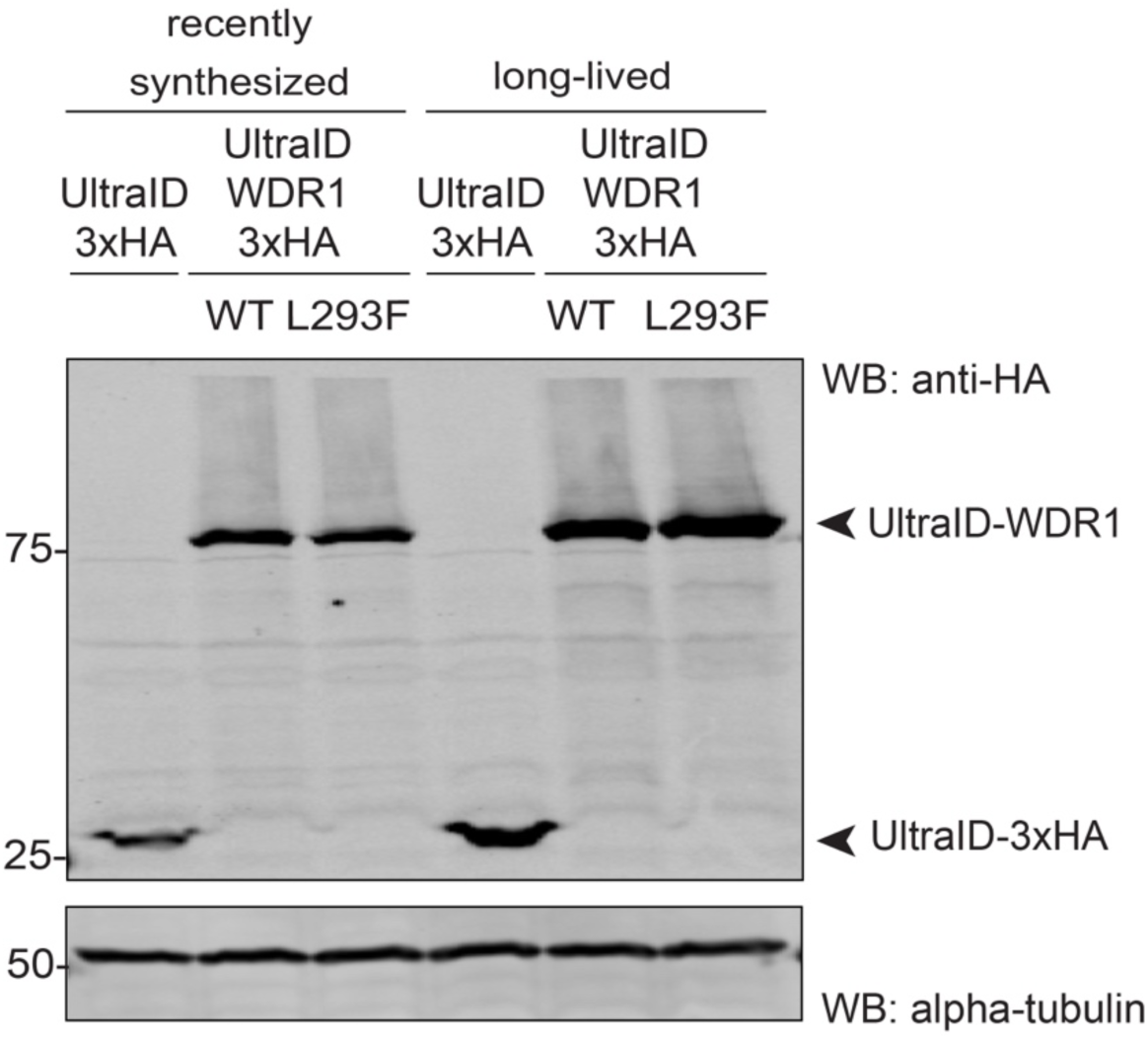
Levels of UltraID-3xHA fused WDR1 WT and L293F are comparable for the reverse-BioID-MS experiments. Western blot analyses of the UltraID-3xHA fused to WDR1 WT and L293F in the recently synthesized (90 mins tetracycline) and the long-lived (4-hour induction and 12-hour chase) conditions.

**Figure S5:**
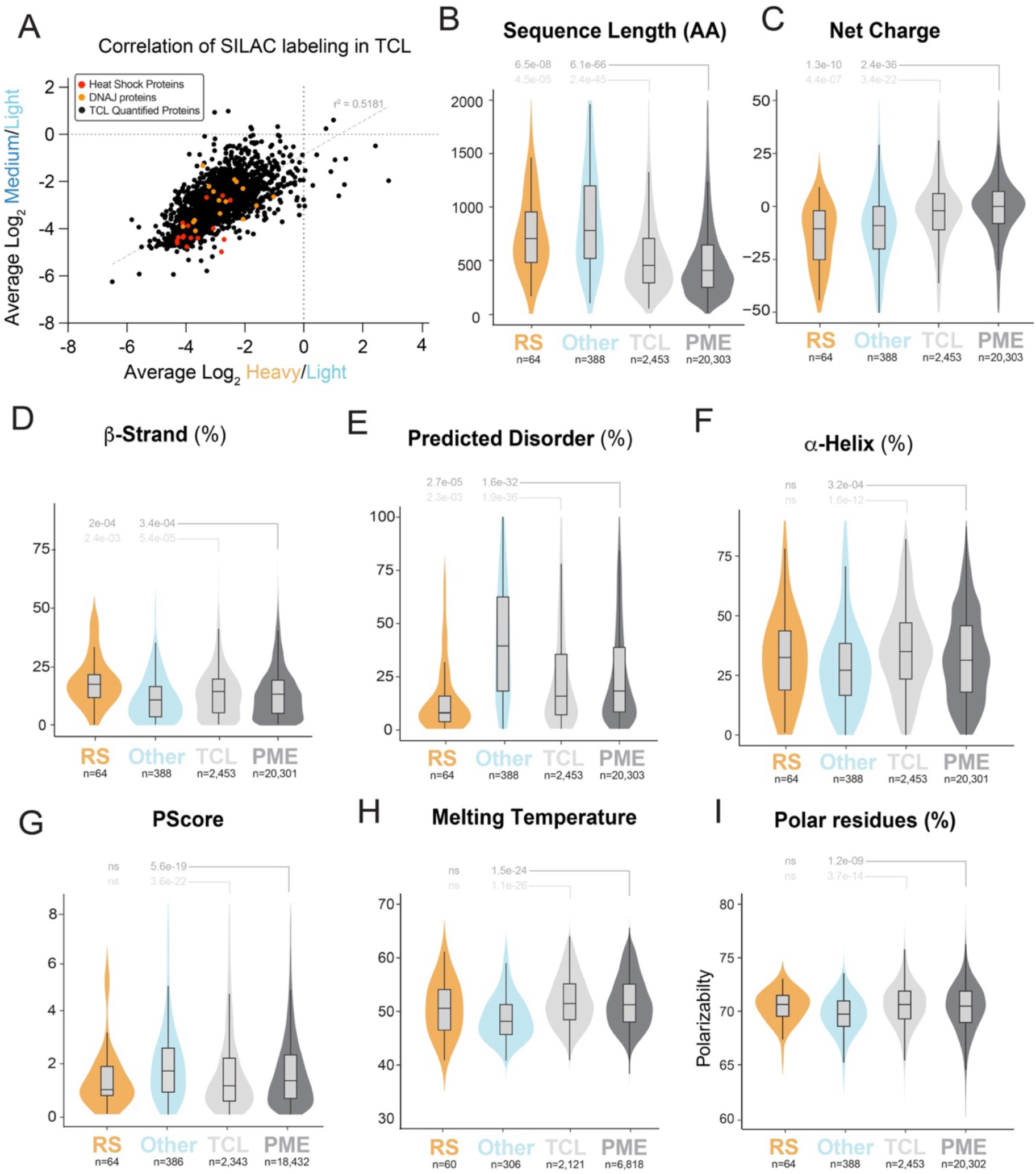
Analysis of recently-synthesized and other DNAJA2 high-confidence interactors during heat stress. **A.** Scatter plot of the average log_2_ H/L and M/L ratios of proteins identified in at least two out of four replicates in the TCL of the heat shock experiment. Each point represents a quantified protein. Quantified DNAJ proteins (orange) and other heat shock proteins (red) are highlighted. **B–I.** Violin plots showing the amino acid (AA) sequence length (B), protein net charge (C), percentage of the residues predicted to form β-strand (D), percentage of residue predicted to be intrinsically disordered (E), the percentage of residue predicted to form alpha-helices (F), protein melting temperatures (G), PScores for pi-pi interactions (H), and the percentage of polar residues (I) among the DNAJA2 high-confidence interactors classified as recently synthesized (RS; n=67) or non-RS (Other, n=388) compared with proteins identified in the TCL or the whole proteome (PME), in the heat shock experiment. Statistical significance was assessed using Wilcoxon rank-sum test with Benjamini Hochberg correction for multiple comparisons.

